# Species richness patterns in Fijian bees are explained by constraints in physiological traits

**DOI:** 10.1101/2024.05.09.593294

**Authors:** Carmen R.B. da Silva, James B. Dorey, Cale S. Matthews, Ben Parslow, Marika Tuiwawa, Julian E. Beaman, Nicholas C. Congedi, Sarah J. Barker, Paris M. Hughes, Rosheen T.E. Blumson, Mark I. Stevens, Michael P. Schwarz, Rosalyn Gloag, Vanessa Kellermann

## Abstract

Determining the ecological and evolutionary mechanisms that underpin patterns of species richness across elevational gradients is a key question in evolutionary ecology, and can help to understand species extinction risk under changing climates. In the tropical montane islands of Fiji, there are 28 species of endemic bee in the subgenus *Lasioglossum* (*Homalictus*), where species richness increases with elevation despite decreasing land surface (habitat) areas. We used a combination of spatially explicit phylogenetic diversity analyses and phylogenetic trait analyses to examine the factors shaping species distributions in these bees. We found that species at higher elevations had lower heat tolerance and desiccation resistance than those at lower elevations, consistent with these traits constraining species’ elevational ranges. We also found high species phylogenetic diversity within mountains, and high phylogenetic signal in species’ heat tolerance and minimum elevational ranges, consistent with these traits being evolutionarily conserved among mountain-top taxa following vicariant (allopatric) speciation. We found no evidence to suggest that interspecific competition is shaping species elevational ranges. In all, our findings indicate that phylogenetic conservatism in physiological traits related to climatic niche, such as heat tolerance, can explain why species richness is highest at mountain tops in this system, with species having tracked their climatic niches over time towards ever higher (cooler and wetter) elevations. Because high elevations in this archipelago are extremely limited (∼2.3% of total land area), only miniscule elevational ‘islands in the sky’ remain into which this diverse, but climate-restricted fauna, can retreat as climates warm.

## Introduction

Determining the processes that shape species geographic ranges and corresponding patterns in species richness across climatic gradients is a key question in evolutionary ecology, especially in the context of climate change. Physiological constraints, evolutionary history, and competition are thought to be some of the key factors that shape species ranges; however, exactly how these factors interact to form range limits has remained a contentious topic for almost two centuries (Forbes 1846, Mayr 1963, Endler 1977, 1982a, Morin and Chuine 2006, Kellermann et al. 2012, Chan et al. 2019, Cadena and Céspedes 2020). Tropical montane island ecosystems are ideal natural laboratories to investigate why patterns in species richness occur across environmental gradients because they often have high levels of endemism and climates change rapidly over limited geographic space (Jankowski et al. 2013).

In montane systems, species richness is often reported to either decrease with increasing elevation, or to have a unimodal relationship with elevation where richness has a mid elevation peak (Rahbek 2005, Hoiss et al. 2012). However, in the tropical montane forests of Fiji, there is an interesting pattern in endemic bee species richness (*Lasioglossum* (*Homalictus)* spp. (Cockerell, 1919); hereafter *Homalictus*), where most species are only found over 800 m above sea level (a.s.l.) (Dorey et al. 2020). Over 20 *Homalictus* species occupy this upper elevational band on one or more peaks within the Fijian archipelago, compared to just four species that can be found at lower elevations (Dorey et al. 2020). This is despite the lower elevations of the islands comprising a far greater area of land than that above 800 m, and the apparent abundance and suitability of floral resources for bees at lower altitude regions.

One possible explanation for the pattern in species richness of Fijian *Homalictus* is that they are broadly adapted to cooler and wetter conditions, and climates have changed faster than has adaptation in their physiological traits, pushing their ranges upslope over time to where the local climates remain cooler and wetter (Ongoma et al. 2021, da Silva et al. 2022). Species geographic ranges are often linked to their physiological traits such as heat tolerance and desiccation resistance (Kellermann et al. 2012a, b, Healy et al. 2019). Indeed, upslope range shifts in response to climate change have been reported in other pollinating insect species, such as butterflies, allowing them to track suitable climatic niches (Rödder et al. 2021, Kerner et al. 2023). On a geological time-scale, climates have cycled repeatedly between warmer-drier and cooler-wetter phases (e.g. Pleistocene climatic cycles), thus today’s patterns in species richness could reflect past rearrangements in species elevational distributions, underpinned by their physiological limits (Patton and Smith 1992).

Physiological traits are in turn influenced by a species evolutionary history. As such, evolutionary history (e.g. speciation processes) can help explain species’ geographic distributions and patterns in species richness (Kellermann et al. 2009, 2012, Cadena and Céspedes 2020). Within tropical montane ecosystems, two key models of speciation have been proposed: gradient/parapatric and vicariant/allopatric speciation (Box 1; Figure 1) (Funk et al. 2016). Both models assume that mountains are species-generating machines, but in different ways (Patton and Smith 1992, Wiens et al. 2019, Cadena and Céspedes 2020, García-Rodríguez et al. 2021), and with different implications for patterns of phylogenetic conservatism of physiological traits across species. The gradient hypothesis is a form of parapatric speciation, where selective pressures (such as climatic variation across an elevational gradient or intraspecific-competition) can drive divergence between populations, leading to speciation across the elevational gradient without physical geographic barriers (Endler 1977, 1982a, b, Rundle and Nosil 2005) (Figure 1A). Thus, the gradient hypothesis assumes that species within mountains should be more closely related to each other than species on different mountains (i.e. species assemblages within mountains should have low 𝛂-phylogenetic diversity) (Figure 1A).

**Figure 1.**
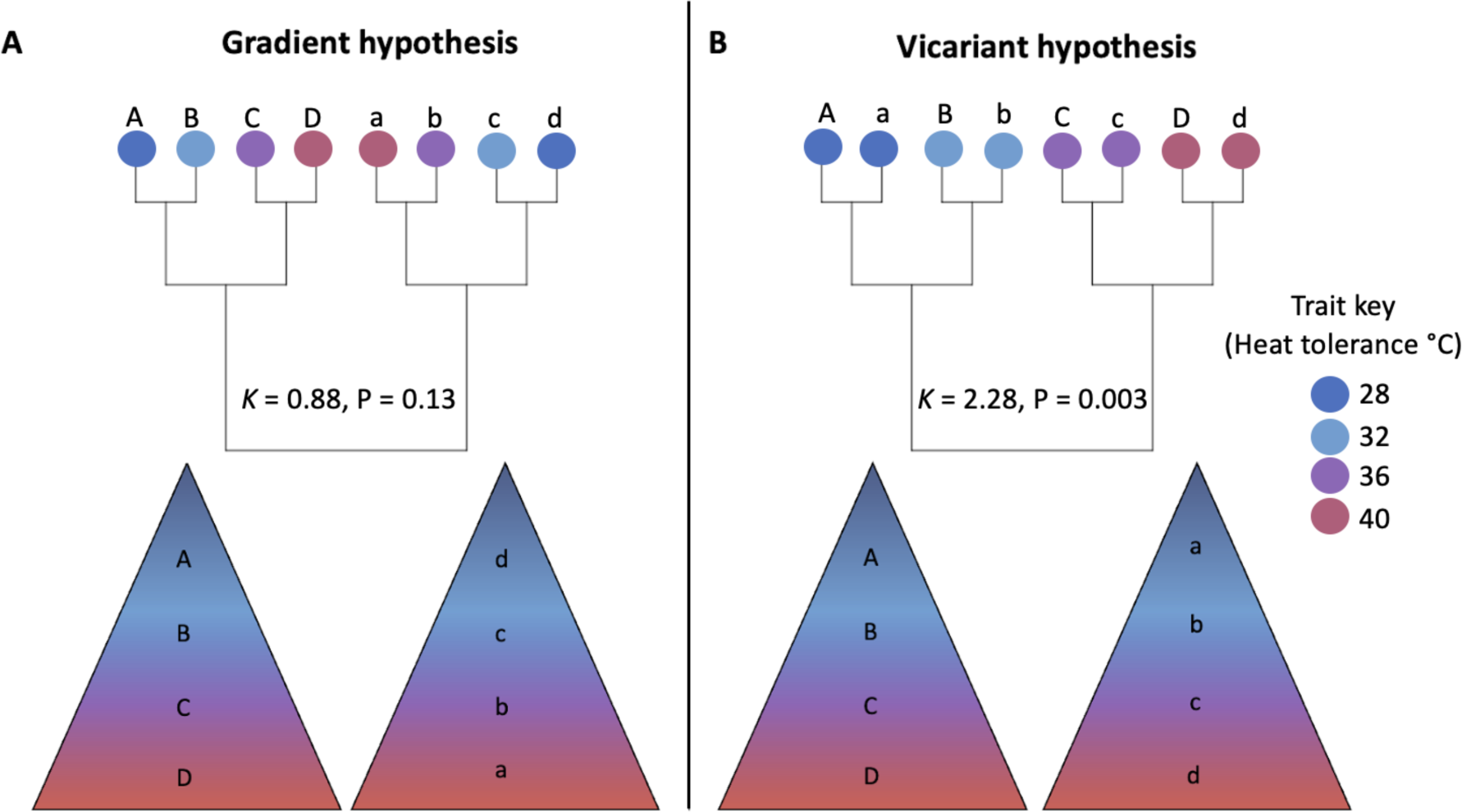
Conceptual figure demonstrating expected phylogenetic and elevational range structure (within and across mountains) for species if they evolved via A) gradient speciation or B) vicariant speciation. Tips of the phylogeny indicate the traits of species (e.g. heat tolerance) and how they correspond to the climates across the elevational gradient (low elevations are warmer and drier than higher elevations). If we simulate phylogenetic signal (represented by Blomberg’s *K* due to low sample sizes) for gradient and vicariant speciation using the above heat tolerance trait values, we find high *K* values for both gradient and vicariant speciation, however, *K* was only statistically significant under the vicariant speciation model. Figure inspired by Patton and Smith 1992. Fluctuating climatic cycles over long geological time periods, and systems with more than the eight species shown in this example are likely to produce phylogenetic patterns that are more complex than those outlined in this conceptual figure.

### Box 1.

Definitions of keywords.

● **Vicariant speciation:** Model of speciation where populations are expected to diversify in allopatry, for example, separation across different mountains.
● **Phylogenetic niche conservatism**: species maintain their ancestral traits (Figure 1B) (Forbes 1846, Mayr and O’Hara 1986, Patton and Smith 1992, Cadena and Céspedes 2020).
● **Gradient speciation:** Model of speciation where species are expected to diversify in parapatry, across elevational gradients (within mountains) due to niche specialisation or competition (Figure 1A) (Endler 1977, 1982a, b, Patton and Smith 1992).
● **Phylogenetic signal**: The propensity for closely related species to have similar traits (Blomberg et al. 2003).
● **𝛂-phylogenetic diversity:** phylogenetic diversity (degree of genetic dissimilarity) within sites (e.g. mountains).

In contrast, the vicariant speciation hypothesis assumes that species evolve via allopatry as a result of population separation by barriers (i.e. mountains or rivers). Under this model, closely related species are not found on the same mountain (Mayr and O’Hara 1986, Patton and Smith 1992, Wiens et al. 2019, Cadena and Céspedes 2020). Rather, species assemblages within mountains will tend to comprise groups of unrelated species (i.e. have high 𝛂-phylogenetic diversity (Figure 1B)). Under gradient speciation, close relatives may show some similarity in traits associated with climate niches (i.e. phylogenetic signal in the trait) within mountain monophyletic groups (Fig 1A), but overall phylogenetic signal within the taxa should be low (Pyron et al. 2015). Meanwhile, vicariant speciation should be associated with high phylogenetic signal in both species climatic niches and related physiological traits (Fig 1B).

In a recent study, Dorey et al. (2020) proposed that phylogenetic niche conservatism can indeed explain why a majority of Fijian *Homalictus* species tend to inhabit high elevation habitats (Dorey et al. 2020). This is because *Homalictus* species show moderate phylogenetic signal in minimum elevational limits (𝜆 = 0.59), consistent with these bees shifting their elevational ranges upslope to track ancestral climatic niches (Dorey et al. 2020). However, whether the apparent conservation of minimum elevation over evolutionary time is underpinned by conservation in physiological traits, or by some other factor, such as competition, remains unknown. For example, if many high-altitude *Homalictus* are in fact physiologically capable of tolerating the heat and desiccation conditions of lower altitudes, then their current distribution is unlikely to be explained by constraints on climate niche. Moreover, the moderate values of phylogenetic signal in minimum elevation detected by (Dorey et al. 2020) could be indicative of either gradient or vicariant speciation (Figure 1) (Pyron et al. 2015), making it challenging to confirm which speciation process is the major driver in this case.

Competition is another major factor that is known to limit species ranges (Terborgh 1971, Terborgh and Weske 1975, Jankowski et al. 2013, Freeman et al. 2022), and thus can contribute to patterns in species richness across space. In closely related species, it has been shown that a species’ range boundary will expand if a competitor species is experimentally removed (Terborgh and Weske 1975). Similarly, in tropical mountain birds, species elevational range breadths are negatively correlated with the number of competitor species found on each mountain (Freeman et al. 2022). Under high competition scenarios, species elevational ranges may thus be truncated compared to their mechanistic niches; that is, they are physiologically tolerant of a larger range of altitudes but are constrained by competition (Jankowski et al. 2013). However, competitive interactions can also be influenced by species physiological tolerances, where certain species will be more competitive under particular environmental conditions (thermal niche partitioning) (Scriven et al. 2016). In the case of Fijian *Homalictus*, climate change and competition might have interacted to produce a pattern where species richness increases with elevation (species shift their ranges to avoid hot and dry climates at lower elevations), but species ranges might be very narrow at high elevations as a result of competitive interactions. In this case, we would predict that for a given mountain, the elevational width of a species’ range decreases as the number of species increases, such that species tend to partition the already space-limited high altitude regions into ever-narrower distributional bands.

In this study, we investigated how species physiological traits, speciation processes, and competition shape the species distributions of Fiji’s endemic *Homalictus* bees. We first generated a robust phylogeny of the clade using a combination of Ultra-Conserved-Elements (UCE) and multilocus datasets. We then paired spatially-explicit phylogenetic diversity and species assembly assessments with assays of two key physiological traits related to climatic niche (heat tolerance and desiccation resistance) to test the hypothesis that phylogenetic conservatism in these traits, following vicariant speciation, explains today’s patterns in species richness with elevation. Finally, we test the prediction that competition also constrains the breadth of species’ elevational ranges. Our data also highlight the vulnerability of endemic Fijian bee species in the context of anthropogenic climate change, a problem that likely faces other tropical biota.

## Methods

### Specimen records

Trends in Fijian *Homalictus* elevational niche conservatism were first explored by Dorey et al. (2020) using collections made between 2011–17 (n = 764). Using the same sampling methods, we undertook additional collections in 2019 (n = 1,203). Collections in 2019 were targeted across elevational gradients and bees were subjected to physiological trait assays (details below), allowing us to examine whether species’ physiological traits might be limiting their elevational ranges to high elevations. The datasets from all years (2011-17 and 2019) were then combined to estimate how species richness changes across elevation, and to evaluate *α*-phylogenetic diversity within mountains.

In total, 1,967 bee collections were made (from 28 *Homalictus* species) across 69 sites across the Fijian archipelago and the entire elevational range (up to 1,324 m a.s.l.). Within Viti Levu, Fiji’s largest island, collections were made on 29 mountains (multiple collection sites within mountains) or coastal areas. Most collections were made by sweep-netting bees from flowering and herbaceous vegetation (native and introduced) or from above the ground near nesting sites. Some additional specimens from the 2011–2017 collections were also collected via passive sampling techniques (Malaise and SLAM traps). At a minimum, for each collection site we recorded latitude, longitude, elevation, date, and collector; the locality data were also checked for accuracy within a week of collection. With the exception of those specimens collected for physiological assays, specimens were collected directly into 98% ethanol and stored at < 5°C. The ethanol was also replaced within a week of collection. All individuals in our dataset were identified to species using CO1 barcoding and/or genitalic morphology as per Dorey et al. (2020).

### Phylogeny construction

To generate a robust phylogeny of the Fijian *Homalictus*, we first generated a UCE tree to constrain the analysis of the multilocus dataset. In total 35 specimens, representing 28 species were included in the analysis. Specimens (n = 18) for the UCE constraint tree were sampled destructively from a single hind leg with DNA extraction conducted at the South Australian Regional Facility for Molecular Ecology and Evolution (SARFMEE) using the Qiagen Gentra Puregene kit (Qiagen Inc., Valencia, CA), following the manufacturer’s protocol. Library preparation and sequencing were conducted at the University of Utah genomics core facility. Extracted DNA was mechanically sheared to a length of ∼600 bp, and individual dual-indexed libraries were generated using llumina TruSeq-style adapters (Glenn et al. 2019). Quantification of adapter-ligated fragments post enrichment was performed via quantitative Polymerase Chain Reaction (qPCR) to ensure capture of UCE loci. Libraries were pooled and sequenced on an Illumina HiSeq 2500 lane. For UCE enrichment we used the Hymenoptera UCE (uce-hym-v2) principal bait set developed initially for ants to enrich 2590-targeted UCE loci across Hymenoptera (Branstetter et al. 2017). For an outgroup, UCE loci were extracted from one genome-enabled species, *Lasioglossum albipes* (ASM34657v1) using the Phyluce Tutorial III (https://phyluce.readthedocs.io/en/latest/tutorials/tutorial-3.html).

#### UCE data processing and alignment

We used the PHYLUCE v1.7.1 pipeline (Faircloth 2016) following tutorial 1 (https://phyluce.readthedocs.io/en/latest/tutorials/tutorial-1.html) to process the raw UCE data. We used the program ILLUMIPROCESSOR (Faircloth 2013), a wrapper around TRIMMOMATIC, to remove adaptor contamination and low-quality reads. We assembled the reads using the wrapper (phyluce_assembly_assemblo_spades) around SPADES genome assembler v3.13.0 (Bankevich et al. 2012) and poorly aligned regions of each alignment were removed using the program GBLOCKS (Castresana 2000) with the following settings: b1=0.5, b2 =0.5, b3=12, b4=7. To determine which probe match setting recovered on average the most loci, six different probe match settings (80–80, 75–80, 70–80, 80–70, 75–70, 70–70) were tested with probe match settings of min-identity 80 and min-coverage 80 (80–80) recovering the highest. The matrices at various levels of completeness were tested individually before selecting the 80% complete matrix as our final dataset. Using the python program spruceup (Borowiec 2016) with default settings and 95% lognormal distribution, a total of 499,000 sites (2.12%) were removed from the data matrix due to potential alignment errors. The summary statistics of the matrix were calculated using AMAS (Borowiec 2016).

#### Phylogenomic Analyses

Partition schemes were determined using the Sliding-Window site Characteristics (SWSC) of site entropy (Tagliacollo and Lanfear 2018), with the best model for nucleotide substitution determined using PartitionFinder v2.1 (Lanfear et al. 2017) using the “rcluster” command (Lanfear et al. 2014) and the “--raxml” command line option to use RaxML v8.0 for calculations (Stamatakis 2006). Phylogenetic inference for each dataset was performed using a concatenated loci maximum likelihood (ML) framework using IQ-TREE v2.1.2 (Minh et al. 2020). For the matrix we specified a partition file allowing individual evolution rates (“-spp” command) and performed 1,000 ultrafast bootstrap replicates (“-bb” command) (Hoang et al. 2018) along with “-bnni” flags to reduce risk of overestimating branch supports.

#### Multilocus data generation and combined tree construction

To build a phylogeny that included species for which we did not have UCE data (n=10 of 28 total species), we extracted three exons that were present in the off-target regions of the UCE contigs. A 375–568 bp fragment of long-wavelength rhodopsin (LWrh; n = 17), a 245–405 bp fragment of Wingless (Wg; n = 18), and a 608 bp fragment of Elongation factor 1-alpha 1 (eEF1a; n = 17). We then concatenated these genes with a 630–658 bp fragment of the mitochondrial cytochrome *c* oxidase subunit I (COI; n = 34) across a total of 35 specimens (duplicates of some species).

The program (phyluce_assembly_match_contigs_to_barcodes) in the PHYLUCE v1.7.1 pipeline was used to extract the loci from the completed pool of contigs. Extracted contigs from the exon capture dataset were checked against the NCBI BLAST database to screen for contamination. Sequences were examined in Geneious v10.2.2 (https://www.geneious.com) for stop codons.

We employed *PartitionFinder* v2.1 (Lanfear et al. 2017) with a greedy, linked and BIC scheme to determine the most appropriate base pair substitution models and combinations for our partitions. We then split each gene into 1^st^, 2^nd^, and 3^rd^ codons positions and found six partitions across the 12 codon positions (Supplementary Table 1). We then used *BEAUti* v2.6.4 to create *BEAST2* run files (Bouckaert et al. 2019). We used a relaxed log normal clock (Lemey et al. 2010) for all partitions except for partition 2 (Table S1), to which we applied a strict clock to ensure convergence. We used a Birth Death tree model and constricted nodes to maintain the tree topology of the UCE constraint tree. We used four heated chains using the package coupledMCMC (Müller and Bouckaert 2020) in *BEAST 2* v2.6.4 (Bouckaert et al. 2019) to adequately explore phylospace and ran each analysis four times. We then combined log and tree files using LogCombiner v2.6.4 (Bouckaert et al. 2019) and checked the traces for convergence (effective sample size >>200) in Tracer v1.7.1 (Rambaut et al. 2018). We then created a maximum clade credibility tree using TreeAnnotator v2.6.4 (Bouckaert et al., 2019) and visualised the tree using FigTree v1.4.4 (Drummond, 2016). Hence, our multi-gene tree matches exactly the topology of our UCE tree. However, the branch lengths and inferred rates will be determined by the multi-gene BEAST2 model.

## Phylogenetic diversity within mountains

We estimated species richness at each sample site, and phylogenetic distance within mountains (𝛼-diversity) using the picante package (Kembel et al. 2010) in the program R (R Development Core Team 2023). We allocated species to mountains and coastal sites by loading all occurrence records into google earth pro (https://www.google.com/earth/versions/#earth-pro) and drawing polygons around each mountain/coastal site (n = 29). These polygons were extracted as a .kml file and converted to a shapefile on MyGeodata converter (https://mygeodata.cloud/converter/kml-to-shp). Each individual bee was assigned a collection site polygon using the *over* function from the *sp* package in R (Pebesma et al. 2012). A community matrix was created to estimate 𝛼-phylogenetic diversity and species richness (Faith 1992) using the picante package at each site and mountain (Kembel et al. 2010). The phylogeny was used to estimate phylogenetic diversity at each site and mountain. We used a second-degree polynomial linear model to examine how phylogenetic diversity and species richness changed across site elevation.

We examined variation in mean phylogenetic pairwise distances between species found within each mountain as an initial step to investigate whether *Homalictus* species evolved in Fiji via gradient or vicariant speciation using the *mpd* function from the *picante* package (Kembel et al. 2010). We estimated the standardised effect sizes of phylogenetic community structure to infer whether phylogenetic distances observed within each mountain are significantly different to null communities generated at random 10,000 times using the *ses.mpd* function. We examined if the standardised effect sizes of mean pairwise phylogenetic distance within mountains was greater or less than 0, and whether they were significantly different to a null species assemblage.

## Phylogenetic signal in physiological traits: Thermal tolerance and desiccation resistance

### Bee sampling

We collected bees for physiological assays on Viti Levu (Fiji’s largest island) during April and September to November in 2019, using the same methods as da Silva et al. (2021). In short, bees were collected by sweep-netting vegetation across multiple altitudinal gradients of 6 to 1,300 m a.s.l. (Supplementary data file 1). We then placed individual bees into large vials (100 mL) with foam lids, allowing aeration immediately upon capture and stored them in a cool, dark container for transport back to the laboratory. We recorded the sex for each specimen through visual assessment (female bees have scopa on their abdomens and 12 antennal segments, whereas males have 13). Most species are cryptic (shiny green) except for *H. hadrander* (blue), *H. groomi* (pink), and *H. achrostus* (black) so we were unable to tell which species were collected and tested during the assays. Instead, species identification was conducted post-assay based on COI barcoding and/or the examination of male genitalia (Dorey et al. 2019). Thus, individual and species sample sizes are different for both assay traits: critical thermal maximum (CT_MAX_) and desiccation resistance (Table S4).

### Physiological trait estimation

We estimated the critical thermal maximum/heat tolerance (CT_MAX_ ; n=9 species), desiccation resistance (n= 6 species), and body mass (n=13) of Fijian bees for a subset of total species collected across the elevational gradient, following the protocols described in da Silva et al. (2021). Briefly, we estimated CT_MAX_ by placing bees into airtight vials within a water bath where the temperature was increased from 25°C at a rate of 0.1°C per minute. CT_MAX_ was scored as the temperature at which individuals ceased movement. We estimated desiccation resistance by placing individual bees into vials with gauze over the top into a desiccation chamber of 5% humidity. We checked bees once an hour until all bees ceased movement; desiccation resistance per bee was recorded as the number of hours elapsed until movement ceased. Following assays, a single hind leg was removed from each individual for COI sequencing. We estimated the dry body mass of each bee using an XP2U Ultra Micro Balance, Mettler Toledo, Victoria, Australia.

### Climate data acquisition

We extracted mean environmental temperature and mean precipitation for each bee’s collection site from WorldClim2 (Fick and Hijmans 2017). We tested the accuracy of the WorldClim2 environmental temperature data which is based on a combination of weather station data, satellite data, and elevational maps by placing HOBO temperature data loggers (www.hobodataloggers.com.au) across the altitudinal gradient at 10 m, 200 m, 700 m, 900 m, and 1,072 m for a period of 2–3 weeks in September-October 2019 (Supplementary Figure 1). On average, the highland was about 5°C cooler and received an additional 100 mm in precipitation each month compared to the lowland region, according to the WorldClim2 data. Based on the HOBO loggers, the mean temperature at the lowland site (Suva, 10 m a.s.l.) during the measurement period was 25.4°C and the mean temperature of the highland site (Tel tower near Nadarivatu, 1072 m) was 20.05°C. Since the HOBO temperature data and WorldClime2 data are well aligned, and because we were unable to measure temperature at each collection site, we used WorldClim2 data to collect environmental data from all sites.

### Phylogenetic trait analyses

We calculated phylogenetic signal in species mean and minimum collection elevation, heat tolerance, desiccation resistance, and body mass using the *phytools* package (Revell 2012). We pooled all data from 2010–2019 to estimate phylogenetic signal in elevation, and pooled all trait data to estimate phylogenetic signal in body mass. Because there are only 28 known species of *Homalictus* in Fiji, and we were unable to measure physiological traits for all of them (as species identification occurred post-trait-assays in most cases) we measured phylogenetic signal using Blomberg’s *K.* An alternative metric, Pagels 𝜆, is known to perform poorly with small sample sizes (Münkemüller et al. 2012, Symonds and Blomberg 2014). However, while the overall species sample sizes are low, we did sample between 21 - 44% of the known *Homalictus* species in Fiji for physiological traits (across different regions of the phylogeny), and 82% for species elevational niches.

To understand whether species physiological traits are associated with constraining species current elevational ranges, we examined the relationship between species collection elevations (mean and minimum) and their physiological tolerances (heat and desiccation resistance) using phylogenetic mixed models with the *rma.mv* function in the metafor package (Viechtbauer 2010) in R (R Core Team 2020). These models incorporate trait standard errors (rather than just trait means) and evolutionary history. Therefore, we used all of the trait data available. Body mass was included as a covariate within all models, so that we could pull apart effects of body size from abiotic factors on species physiological traits. We extracted the mean maximum environmental temperature and precipitation of the driest month from each species’ collection locations. However, these climatic variables are highly correlated with collection elevation, and models that included elevation and the climatic factors had variance inflation factors over 2.5. Therefore, we excluded the climatic factors from our models to avoid issues associated with multicollinearity, but kept collection elevation as a predictor variable (Johnston et al. 2018). We found that a model that included elevation as a second degree polynomial better explained variation in heat tolerance, but, the relationship between desiccation resistance and elevation was linear, so no polynomial functions were added to the desiccation model. Significance of fixed effects was tested using likelihood ratio tests. We used the packages *phytools* (Revell 2012) and *ggplot2* v3.30 to produce data figures (Wickham 2016).

## Competition

To examine whether competition contributed to patterns in species’ elevational ranges, we assessed the relationship between median elevational range across all species on each mountain and species richness, following the methods of Freeman et al. (2022). We first calculated each species’ elevational range on each mountain and then estimated the median elevational range that species occupy on each mountain. We used median elevational range, rather than mean elevational range in this analysis because one species, *Homalictus fijiensis*, has a very large elevational range across all mountains, but is most abundant in the lowlands, while most other species have very narrow ranges. Using the median value thus better captures the relationship between richness and elevational ranges in our dataset. We excluded coastal sites from this analysis. Because the data were not normally distributed, we log-transformed median species elevational range and species richness and then used a Pearson’s product-moment correlation. If competition was a key factor shaping the elevational breadth of species’ ranges, then we expected that species elevational ranges would decrease as species richness increased. If competition had little or no impact on the elevational breadth across which a species occurred, then we expected no relationship between species richness and species elevational ranges.

## Results

Our data set comprised 1,967 *Homalictus* specimens from across the Fijian archipelago collected between 2011 and 2019. We recovered 28 *Homalictus* species across the entire elevation gradient in Fiji (sample sizes for each species, and summary elevational distributions are given in Table S1).

### Species richness and phylogenetic diversity

We found that richness increased with elevation across sites (F_2,55_ = 38.21, *p* < 0.001; Figure 2). In particular, there was an increase in species richness beginning from about 600 m asl, despite the fact that total land area per elevational band dramatically decreases from this altitude (Figure 2). Similarly, phylogenetic diversity (an alternative to measuring species richness and a metric that incorporates genetic variation) also increased with elevation across collection sites (F_2,55_ = 37.77, *p* < 0.001).

**Figure 2.**
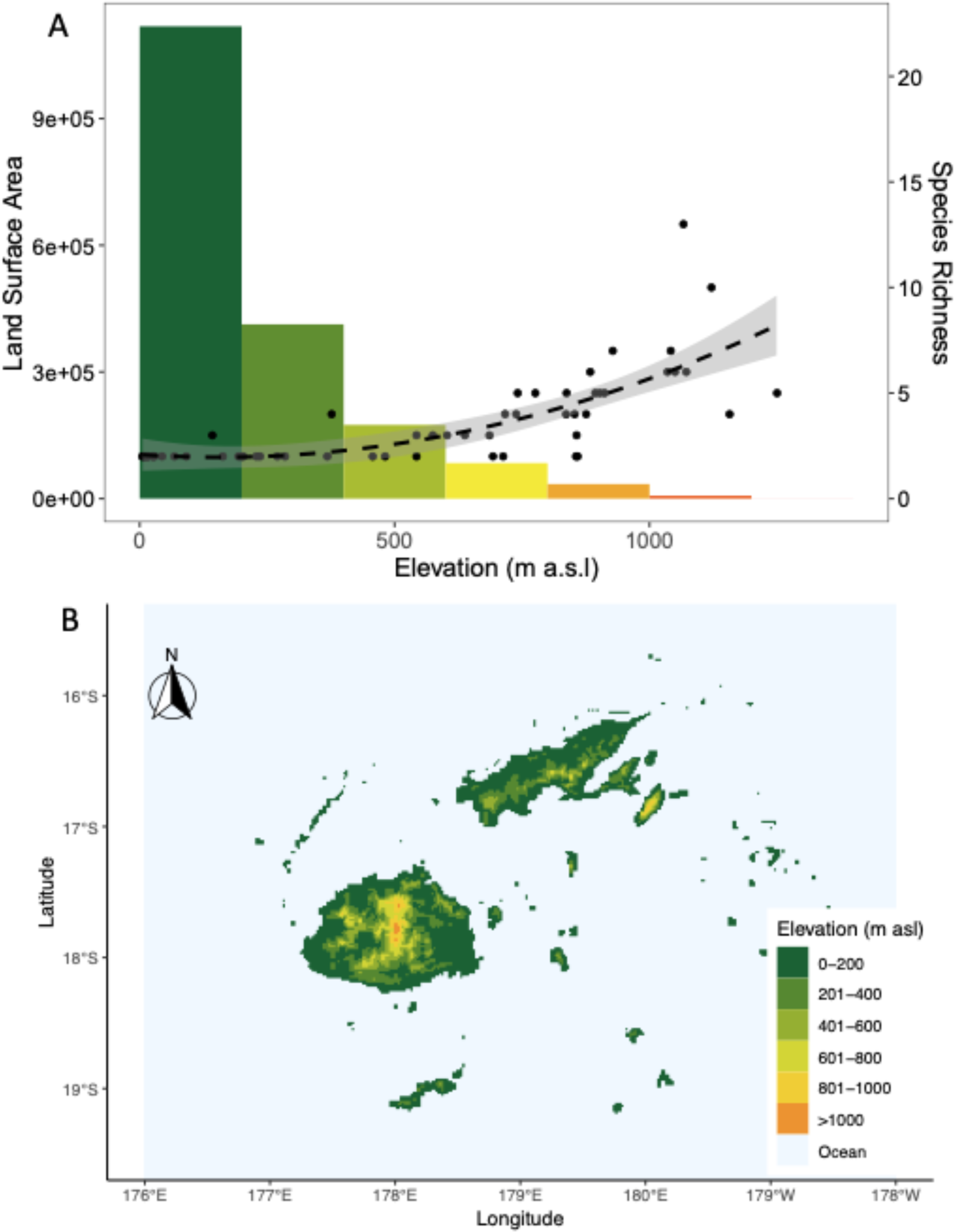
(A) Land surface areas of Fiji in 200 m asl bands (histogram) and *Homalictus* species richness (number of species) across collection site elevation (B) Map of Fiji showing a digital elevation model in 200 m a.s.l. bands, where colours correspond to top panel. Figue S3 shows the elevational ranges of each species explicitly.

As a first step towards understanding the role of evolutionary history on current species distributions, we assessed whether species on the same mountain (combining multiple sites within mountains) were phylogenetically clustered by measuring α-phylogenetic diversity. Mountains with positive standardised effect sizes of mean pairwise phylogenetic distance and high quantiles (mpd.obs.p > 0.95) indicate high phylogenetic diversity (i.e low relatedness of species assemblages within mountains), consistent with vicariant speciation. Whereas negative standardised effect sizes of mean pairwise phylogenetic distance and low quantiles (mpd.obs.p < 0.05) indicates phylogenetic clustering (i.e. closely related species are found on the same mountain) (Kembel et al. 2010), consistent with parapatric or sympatric speciation. We found that the standardised effect sizes of mean pairwise phylogenetic diversity within each mountain with species richness greater than 1 (n=16) were either above 0 with high quantiles (high phylogenetic diversity), or less than 0 but with quantiles greater than 0.05 (low phylogenetic diversity, but not significantly different to a randomly generated null assemblage), suggesting that species assemblages within all mountains are not more closely related than we would expect at random (Table S2; Figure 3). These estimates of phylogenetic diversity are therefore most consistent with *Homalictus* species in Fiji having speciated via vicariance.

**Figure 3.**
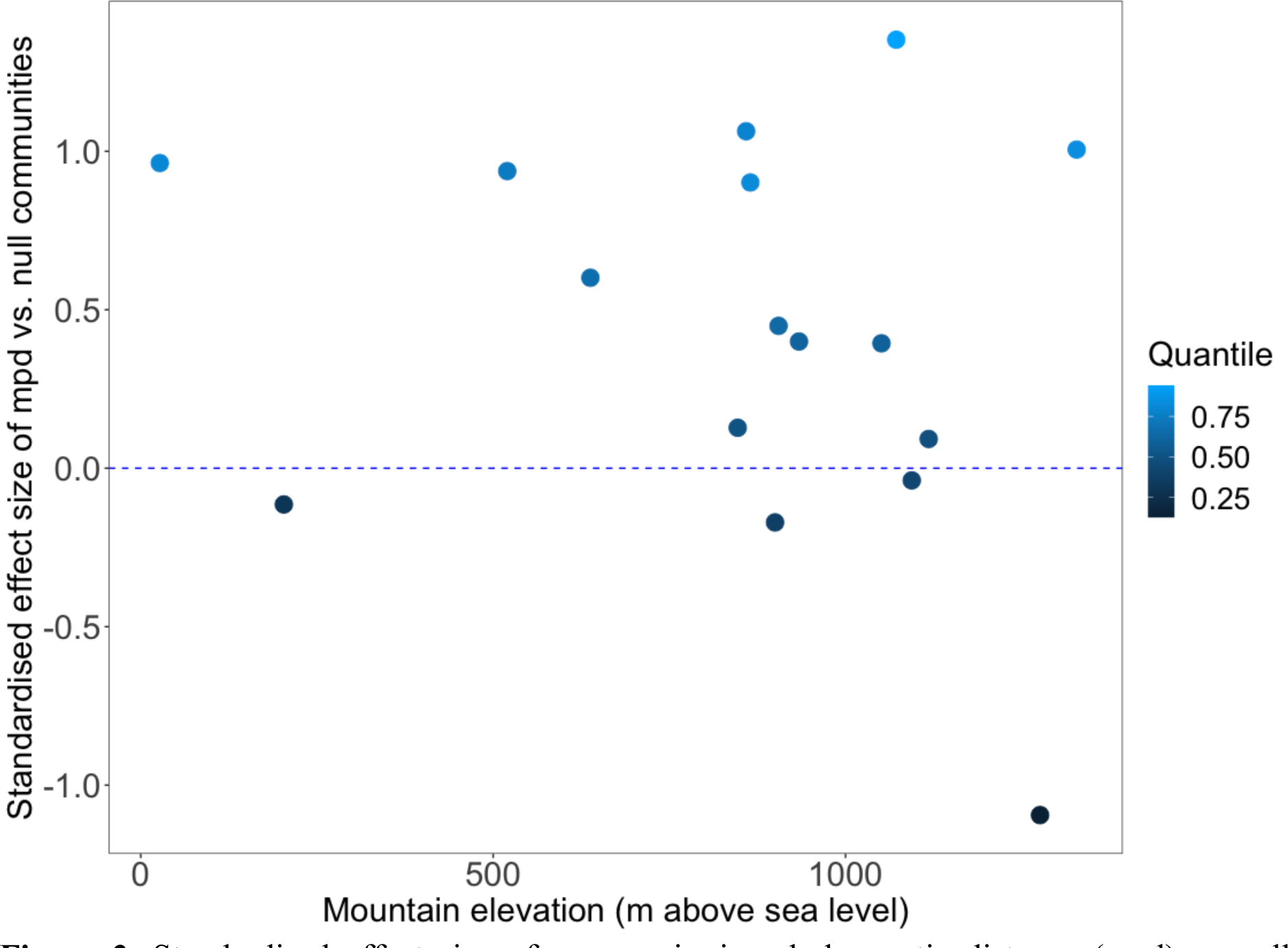
Standardised effect size of mean pairwise phylogenetic distance (mpd) vs null communities across mountains of different elevation. While some points have standardised effect sizes of mpd below 0 (indicating phylogenetic clustering), their quantiles are greater than 0.05, suggesting that they are not more closely related than we would expect compared to a random community assemblage (see Table S2). Only sites with more than one species are shown in this graph (n=16) because phylogenetic distance can not be calculated with less than 2 species (only *H. fijiensis* was collected at some coastal sites). ɑ-phylogenetic diversity data is included within additional dataset 2.

### Phylogenetic signal in physiological traits

As a second step towards understanding how evolutionary history might have shaped species current geographic ranges, we assessed phylogenetic signal in species physiological traits and elevational ranges.

We found strong phylogenetic signal in species minimum collection elevation (Table 2; Figure 4), heat tolerance (Table 2; Figure 5A), and body mass (Table 2). We did not, however, find evidence of phylogenetic signal in mean collection elevation or desiccation resistance (Table 2; Figure 5B). Elevation was the only variable that was collected for all species with sequence data, therefore, we show the full phylogeny with minimum collection elevation mapped across the phylogeny (species with sample sizes of 1 are not shown; Figure 4). To compare estimates of phylogenetic signal in minimum range elevation to Dorey et al. (2020) with our expanded dataset we also calculated Pagel’s 𝜆 for minimum range elevation (Figure 4).

**Table 2.** Phylogenetic signal (Blomberg’s *Κ*) for niche-related traits in endemic Fijian *Homalictus* species. Each estimate of *K* is shown with an associated *p* which indicates whether phylogenetic signal estimates are significantly different from what we would expect at random for each measure of phylogenetic signal. Phylogenetic signal was only calculated for species with sample sizes greater than 1 for elevation (minimum and mean) and desiccation resistance. Exclusion of species with sample sizes of less than 1 did not significantly impact estimates of phylogenetic signal for all other traits so all species were included in these estimates of phylogenetic signal (i.e. only 1 individual of *H.* new species *1* was collected for heat tolerance). *indicates whether the phylogenetic signal is statistically significant.

| Trait | $K$ | $p$ | n |
| --- | --- | --- | --- |
| Mean elevation | 0.66 | 0.131 | 25 |
| Minimum elevation | 1.01 | 0.003* | 25 |
| Heat tolerance | 1.3 | 0.02* | 9 |
| Desiccation resistance | 0.72 | 0.14 | 6 |
| Body mass | 0.94 | 0.01* | 13 |

**Figure 4.**
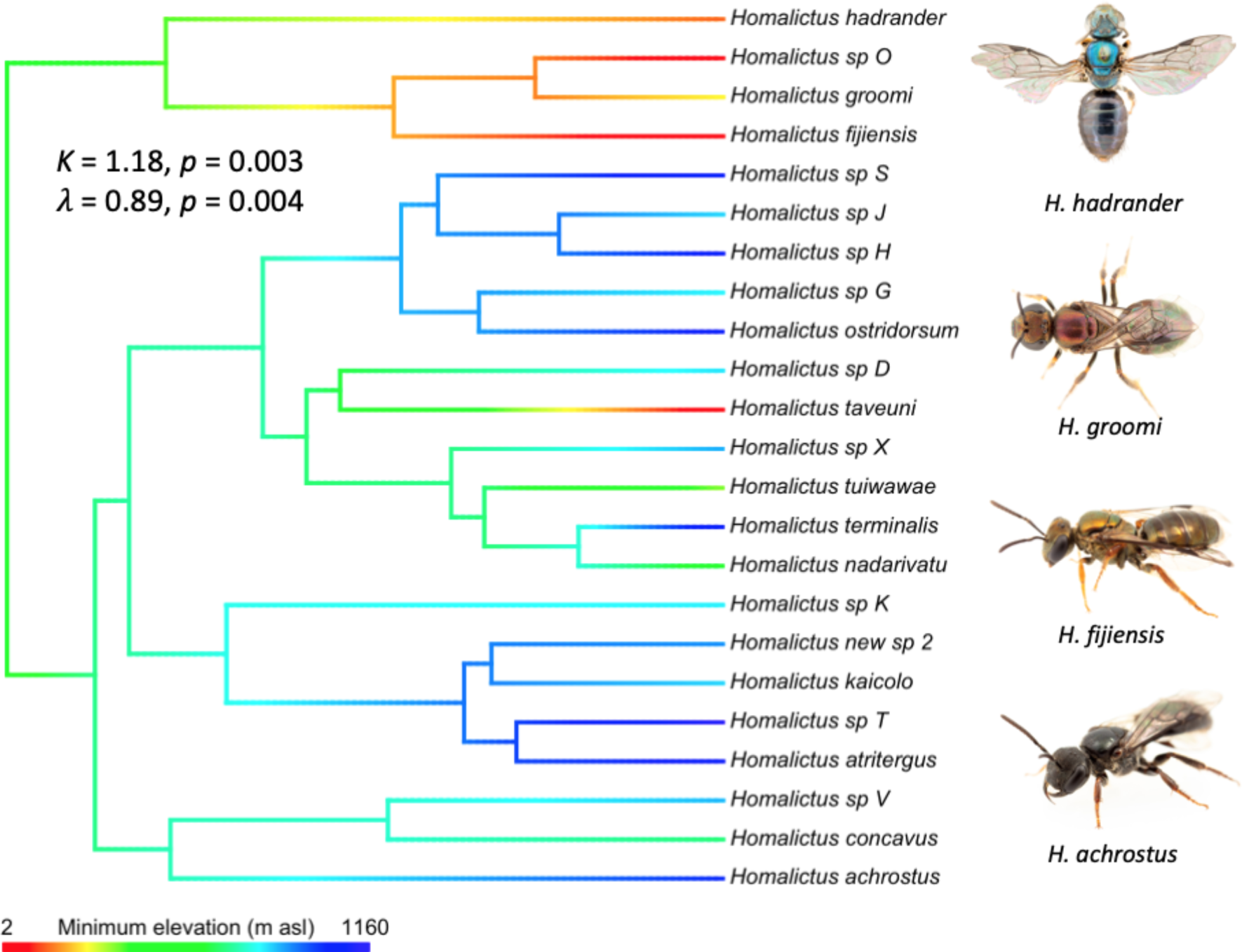
Phylogeny of endemic Fijian *Homalictus* generated using ultraconserved elements and multilocus data. Minimum collection elevation is mapped across the branches to visualise phylogenetic signal in elevational niche. Five of the 28 total species were excluded from this figure due to having a sample size of 1 (*H. taveuni*, sp. I, sp. F, sp. W, and new sp. 1); see full phylogeny in Supplementary Figure 2). Photos by JBD.

**Figure 5.**
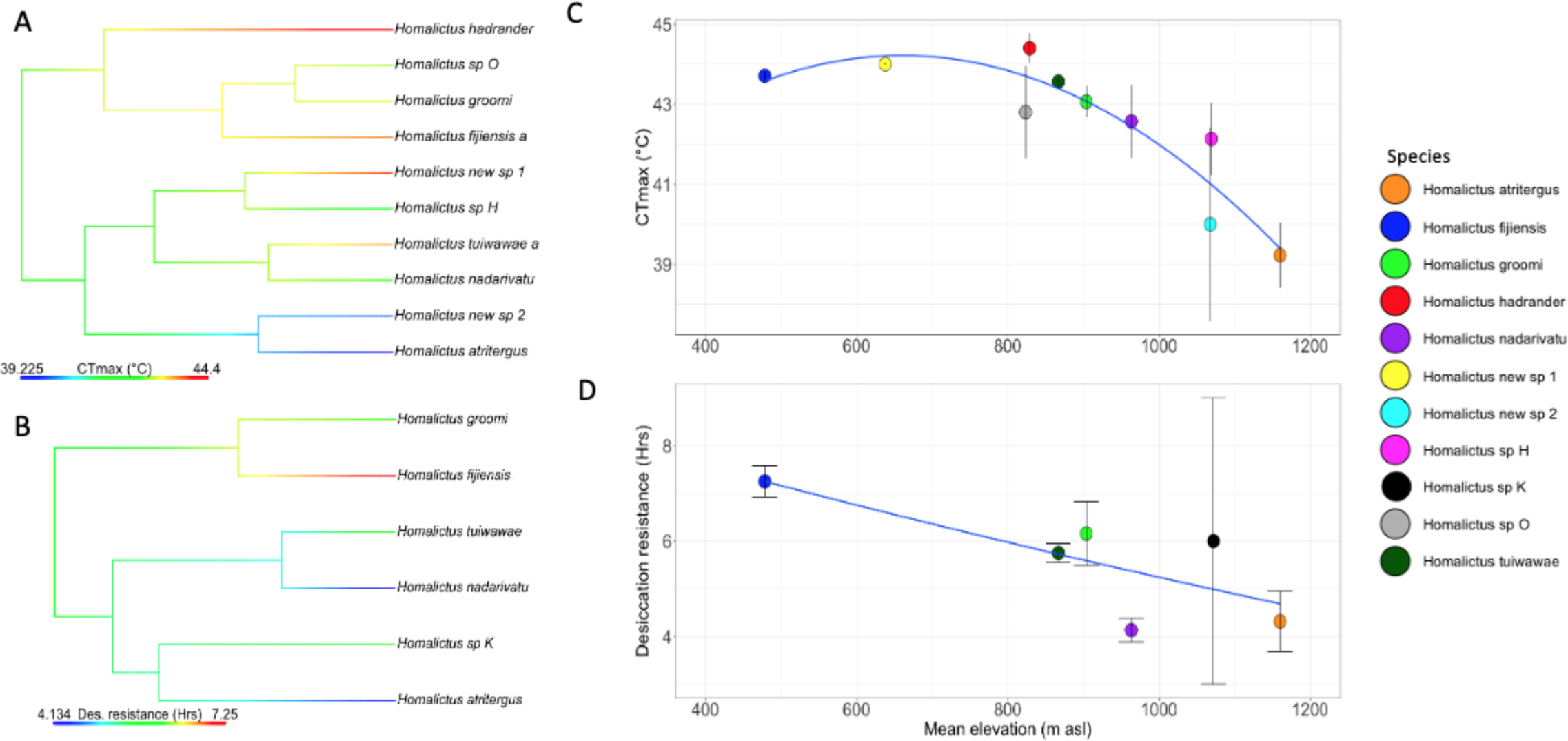
A) Heat tolerance (CT_max_) and B) desiccation resistance mapped across the Fijian Homalictus phylogeny. C) Mean CT_max_ (°C) (± se) and D) mean desiccation resistance (Hrs) (± se) across species mean collection elevations (m asl).

To determine if species physiological traits could be restraining their current geographic ranges, and resulting patterns in species richness across space, we examined correlations between physiological traits and elevation. Our phylogenetic mixed models show that mean collection elevation explains significant variation in species heat tolerance (ꭓ^2^ = 12.85, *p* = 0.002) (Figure 5C), but body mass did not explain variation in heat tolerance across species (ꭓ^2^ = 3.02, *p* = 0.082). Species that are found at high elevations (which are cooler on average) were less heat tolerant than species found at lower and warmer elevations (Figure 5C).

Similarly to heat tolerance, variation in desiccation resistance is explained by mean collection elevation (ꭓ^2^ = 7.05, *p* = 0.029) (Figure 5D), but not mean body mass (ꭓ^2^ = 4.29, *p* = 0.12). Variation in body mass was not explained by species’ mean collection elevations (ꭓ^2^ = 0.86, *p* = 0.36).

### Competition

We did not find any evidence that competition within mountains was affecting species median elevational ranges (rho = 0.25, *p* = 0.32) (Figure 6). That is, we found no relationship between the number of species on a mountain and the breadth of species’ elevational ranges (whereas under competition, we would predict high-richness mountains to result in narrower ranges per species).

**Figure 6.**
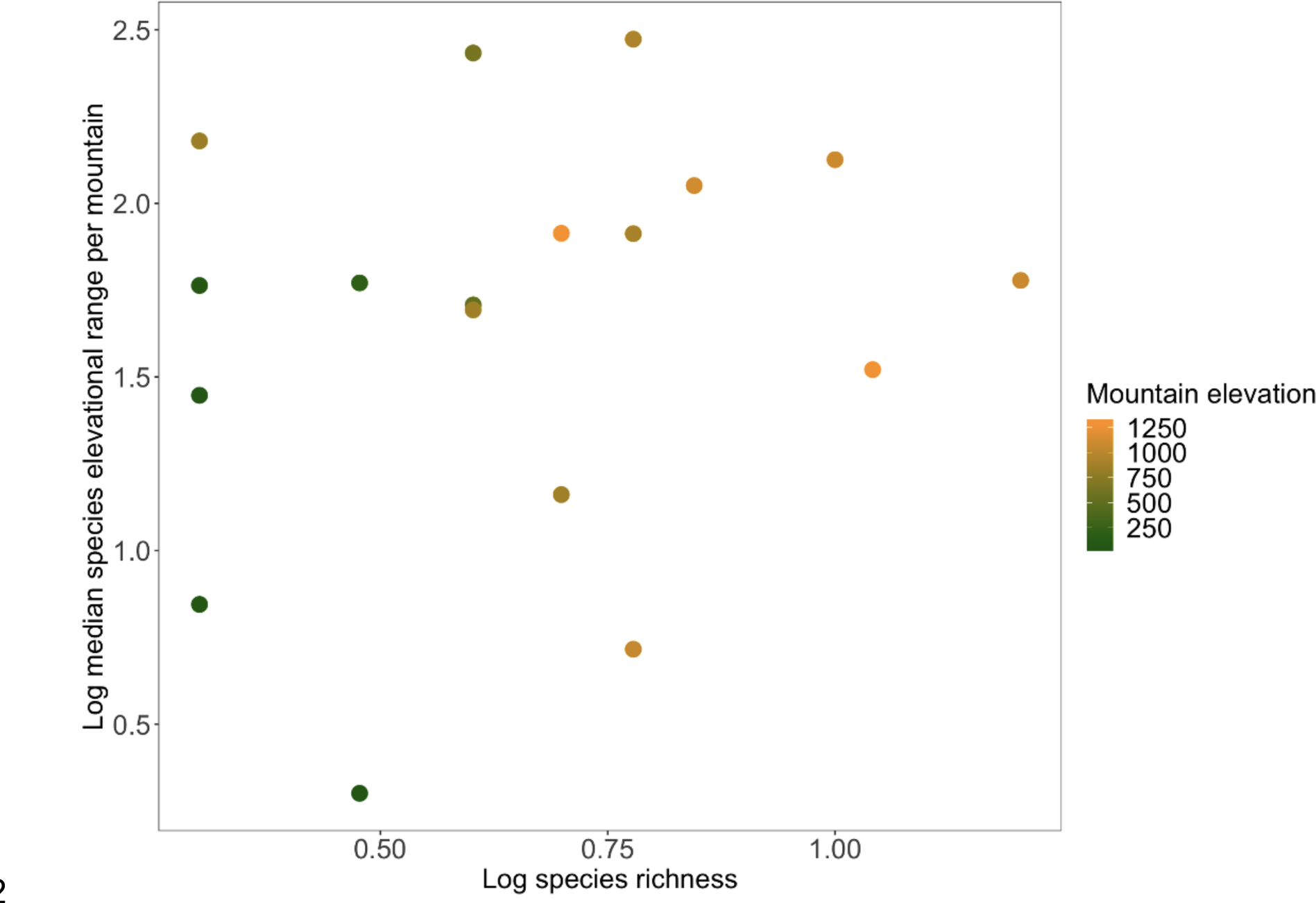
Log-transformed median elevational range of species on each mountain depending on the log species richness on each mountain. Dots are coloured by mountain elevation.

## Discussion

Understanding the processes that shape species ranges and spatial patterns in species richness remains a key goal for ecologists, and is important for predicting how species ranges and assemblages will shift with further anthropogenic climate change (Morin and Chuine 2006, Kellermann et al. 2012, Diamond 2018). We found that for *Homalictus* bees in Fiji: (i) species richness increases with elevation, (ii) closely related species were found on different mountain tops, consistent with vicariant (allopatric) speciation, (iii) closely related species also had similar heat tolerances and minimum elevational ranges, providing support for phylogenetic niche conservatism in these traits, (iv) species’ heat tolerance and desiccation resistance decreased as elevational range increased, consistent with these traits limiting species’ ranges, and (v) the breadth of species’ elevational range did not decrease when more species were present on a mountain, suggesting competition with other *Homalictus* was not a major driver of species’ ranges. Given the evidence for conserved thermal physiology and elevational niches over evolutionary time, it seems likely that as climates continue to warm, Fijian *Homalictus* will track their climatic niches to ever higher elevations, where possible.

Mountains are known to be species generating machines due to the varied climatic niches and physical barriers they present (García-Rodríguez et al. 2021). However, the relationship between elevation and species richness varies across systems. While we found that species richness in Fijian *Homalictus* increases with elevation, many other systems show the opposite trend, with decreasing species richness with elevation (e.g. Rahbek 2005, Hoiss et al. 2012), including other Hymenoptera in Fiji. For example, Fijian ants have their highest species richness at lower elevations (Economo and Sarnat 2012). Owing to the extreme thermal tolerances of ants, where even high elevation populations (1,300 m a.s.l.) can have very high thermal tolerances (LT50) of 57 °C (Villalta et al. 2020), perhaps their geographic distribution patterns have not been impacted by changing climatic cycles in the same way as the more thermally-sensitive *Homalictus* bees. Speculatively, the opposing pattern in species richness across elevation in Fijian ants compared to that observed in Fijian *Homalictus* bees suggests that different processes underlie speciation in these two related insect groups in Fiji.

Our results indicate that *Homalictus* species in Fiji most likely speciated in allopatry following the repeated isolation of ancestral populations on different mountaintops. Phylogenetic diversity within each of the mountain/coastal regions we sampled was high, and we found no significant phylogenetic clustering of closely related species within mountains. Other studies in montane ecosystems have similarly found that species that replace each other along elevational gradients are not closely related, including in mammals (Patton and Smith 1992) and birds (Cadena et al. 2012, Caro et al. 2013, Cadena and Céspedes 2020). The complex topology of mountains, and thus the high frequency of separation by barriers, is thought to drive vicariant (allopatric) speciation at rapid rates, making mountains ‘biodiversity pumps’ and explaining why species richness is often so high in the world’s vegetated mountainous regions (Moritz et al. 2000, Kozak and Wiens 2006, García-Rodríguez et al. 2021). Yet there is also empirical support in other studies for gradient (parapatric) speciation, in which populations diverge due to local adaptation (Chapman et al. 2013, Funk et al. 2016). It may be that often both speciation processes may play a role over different time scales or on different mountains within a taxon’s range. This may be the case also for *Homalictus*, given that it is challenging to estimate the mechanisms that have driven divergence between all clades present in Fiji today. For example, an absence in phylogenetic clustering within mountains could also be influenced by high dispersal during cool climate cycles. However, given that we also observed very high phylogenetic signal in both minimum elevational range and heat tolerance, vicariant speciation coupled with phylogenetic niche conservatism is a more parsimonious model to explain Fijian *Homalictus* present day geographic ranges.

Our data are consistent with climate-related physiological traits constraining the elevational niches of Fijian *Homalictus*. Species collected at high elevations tended to have lower heat tolerances and were less desiccation resistant than those found at lower elevations. High elevation regions in Fiji tend to be ∼5°C cooler, and receive about 100 mm more rainfall each month, than the lowland regions (da Silva et al. 2022). Thus trends in these traits reflect changes in climate across altitudinal space. Studies that link variation in heat tolerance to species ranges and evolutionary history remain surprisingly rare; however, research on both *Drosophila* and freshwater fishes have highlighted that heat tolerance can be evolutionary constrained and linked to species geographic ranges (Kellermann et al. 2009, 2012, Comte and Olden 2017). Across broad taxonomic groups of terrestrial ectotherms, however, heat tolerance is not correlated with latitude or elevation (i.e. species ranges) (Sunday et al. 2011, 2019). In addition, climate origin of ancestry does not shape heat tolerance, suggesting that heat tolerance is not conserved across broad taxonomic groups on a global scale (Bennett et al. 2021). Why we might observe phylogenetic signal and tight associations between species heat tolerances and their geographic ranges within closely related species (i.e. within genera, such as the *Homalictus* in this study) but not across diverse taxa remains an open question.

We did not find phylogenetic signal in desiccation resistance for Fiji’s *Homalictus*. This might reflect low power in our study to detect signal in desiccation resistance; only six species were tested for this trait, although these were spread across the phylogeny and represent around one fifth of the Fijian *Homalictus* phylogeny. Alternatively, it may be that desiccation resistance is in fact more evolutionarily labile than heat tolerance. That is, bees may readily evolve differences in desiccation resistance depending on their local environmental conditions, or they may be more likely to respond to changes in climate via phenotypic plasticity (i.e. developmental plasticity or reversible acclimation). For example, in a laboratory experiment, desiccation resistance was found to evolve rapidly in a polyphagous fly (Tejeda et al. 2016), and fruit flies have the capacity to plastically alter their desiccation physiology (i.e. rates of evaporative water loss) (Bosua et al. 2023). Overall, *Homalictus* species in Fiji tended to be fairly intolerant of desiccating conditions, with all species showing mean desiccation resistances in our assays of less than 8 hours. This limited desiccation tolerance highlights the deep tropical origin of *Homalictus* bees (Ibalim et al. 2020). In comparison, two common invasive stem nesting bees with broad geographic distributions, which are common in the lowland region of Fiji, *Braunsapis puangensis* (Cockerell, 1929) and *Ceratina dentipes* Friese, 1914, have mean desiccation resistances of 20 and 30 hours, respectively (da Silva et al. 2021).

The close correlation between species mean elevations and their physiological traits is consistent with the idea that high-altitude *Homalictus* (a majority of the *Homalictus* species) are unable to tolerate the warmer and drier conditions found at lower elevations. However, most *Homalictus* species, even those at mountain tops, had heat tolerances higher than 40°C, which is warmer than the highest air temperature on record in Fiji (35°C recorded in January 2013 by Nausori weather station). High temperatures are known to have sublethal effects on an individual’s survival and fitness before they reach their critical upper thermal limits (da Silva et al. 2020, van Heerwaarden and Sgrò 2021), and these sublethal effects may ultimately determine species’ elevational ranges. Additionally, the behaviour of ground-nesting bees (such as *Homalictus*) can be strongly impacted by ground surface temperatures (Antoine and Forrest 2021), which can far-exceed air temperatures (Kearney 2019), potentially altering the frequency at which bees can enter and leave their nests, thus impacting their elevational distributions. Finally, limited desiccation resistances are also likely playing a major role in restricting many species’ ranges to high elevations, which are much wetter than the lowlands. High environmental temperatures are associated with increasing vapour pressure deficits (drying power of the air) (da Silva et al. 2022), so even though species might have high thermal tolerances, if they are not desiccation tolerant they will not be able to inhabit warm environments if humidity is not high.

We did not find any evidence suggesting that competition has shaped species elevational niches or species richness in Fiji. This finding is in contrast to several previous studies which reveal competition curtailing species elevational limits (Terborgh 1971, Terborgh and Weske 1975, Freeman et al. 2022). *Homalictus* species are known to be generalist foragers, making use of a wide variety of floral resources, which may minimize resource competition in these bees (Crichton et al. 2019, Draper et al. 2021). Furthermore, food resources and ground nesting sites are likely to be plentiful in Fiji’s highly productive highland regions (due to high precipitation), further reducing interspecific competition within the genus. In other systems, where resources are potentially more limited, climate has been proposed to impact the strength of competition, where harsher climates are expected to strengthen competition (Chan et al. 2019). Perhaps this pattern is less pronounced therefore in tropical ecosystems, which generally lack harsh extreme temperatures or very low humidities.

### Species risk under further climate change

As climates continue to warm with anthropogenic climate change, *Homalictus* species are likely to be pushed to even higher elevations to track their climatic niches. Alarmingly, 90% of the species in Fiji already inhabit elevations higher than 800 m a.s.l., and with the tallest mountain in Fiji, Tomanivi, at 1,324 m a.s.l., species will soon have no further elevational refugia to which to retreat. Behavioural thermoregulation, such as hiding in deep ground nests during unfavourable climatic conditions, might provide some respite from extreme temperatures (da Silva et al. 2021). However, reduced foraging activity outside the nest on high temperature days could also have implications on brood provisioning and pollination (Jaboor et al. 2022). Evolutionary change is the only viable long-term solution to survival with climate change when range shifts are not possible, especially as plastic responses are often small (Gunderson and Stillman 2015). Unfortunately, the high phylogenetic signal in species minimum elevational ranges and heat tolerances suggests that there are likely evolutionary constraints impeding *Homalictus* from rapidly adapting to a changing climate (Dorey et al. 2020, Martin et al. 2023).

## Conclusion

We found that vicariant speciation and species physiological traits (heat tolerance and desiccation resistance) have shaped *Homalictus* species present day geographic ranges in Fiji, and coinciding patterns in species richness across an elevational gradient. A majority of Fijian *Homalictus* species are only found at elevations above 800 m a.s.l. These ‘sky island’ species are likely to be particularly vulnerable to further climate change as they have limited higher elevation refugia to retreat to, and they are likely to have a limited capacity to adapt to the warmer and drier climates found at lower elevations (implied by phylogenetic niche conservatism). Tropical ectotherms tend to be under-represented in studies of climate change vulnerability assessments (White et al. 2021), even though they are expected to be the least resilient to warming climates (Deutsch et al. 2008). Future research and conservation efforts should prioritise tropical ectotherms before we lose key species.

## Acknowledgements

We would like to thank people from the highland villages of Navai and Nadarivatu for access to high elevation collection sites and for welcoming us into their villages. We would like to thank Dr. Lesley Alton (Monash University) for assisting bee collections in April 2019 and Dr. Stephen Galvin (University of the South Pacific) for logistical support in Fiji. We thank Dr. Emily Sadler for conducting the library preparation and ultraconserved element sequencing.

## Author Contributions Statement

CRBdS, RG, MPS, and VK conceptualised the research project. CRBdS, VK, JBD, JEB, SJB, NCC, PMH, and RTEB conducted field work and physiological trait assays in Fiji. MT provided crucial logistical support in Fiji. MIS and MPS funded the genetic sequencing and conducted field collections. JBD and BP made the Fijian *Lasioglossum (Homalictus)* phylogeny. CRBdS conducted the statistical analyses and wrote the initial manuscript draft. All authors edited the draft.

