## Supplementary material for "Species richness patterns in Fijian bees are explained by constraints in physiological traits"

**Supplementary Tables and Figures**

**Table S1**: The gene partitions and best substitution model as chosen by *PartitionFinder* version 2.1 (Lanfear et al., 2016). The genes are a 630-658 bp fragment of the mitochondrial cytochrome *c* oxidase subunit I (COI), a 375-568 bp fragment of long-wavelength rhodopsin (LWrh), a 245-405 bp fragment of Wingless (Wg), and a 608 bp fragment of Elongation factor 1-alpha 1 (eEF1a).

| Partition number | Gene codons | Best model |
|---|---|---|
| 1 | EF1a-2$^{nd}$ and COI-2$^{nd}$ | K80+I |
| 2 | Opsin-1$^{st}$, 2$^{nd}$, 3$^{rd}$, and EF1a-3$^{rd}$ | K80+I |
| 3 | Wg-1$^{st}$, 2$^{nd}$, and EF1a-1$^{st}$ | JC |
| 4 | Wg-3$^{rd}$ | HKY+I+Γ |
| 5 | COI-1$^{st}$ | GTR+Γ |
| 6 | COI-3$^{rd}$ | F81 |

**Table S2.** Summary of species mean collection elevations and sample sizes.

| **Species** | **Mean elevation** | **n** |
|---|---|---|
| *Homalictus sp.* D | 885.7 | 3 |
| *Homalictus sp.* F | 1089.0 | 1 |
| *Homalictus sp.* G | 890.0 | 2 |
| *Homalictus sp.* H | 1069.3 | 3 |
| *Homalictus sp.* I | 27.0 | 1 |
| *Homalictus sp.* J | 898.5 | 2 |
| *Homalictus sp.* K | 1029.5 | 6 |
| *Homalictus sp.* kaicolo | 1036.8 | 4 |
| *Homalictus sp.* O | 823.5 | 19 |
| *Homalictus sp.* S | 1118.0 | 36 |
| *Homalictus sp.* T | 1226.9 | 8 |
| *Homalictus sp.* V | 905.4 | 5 |
| *Homalictus sp.* W | 1070.0 | 1 |
| *Homalictus sp.* X | 1226.5 | 36 |
| *Homalictus achrostus* | 1042.0 | 2 |
| *Homalictus atritergus* | 1160.2 | 48 |
| *Homalictus concavus* | 800.7 | 14 |
| *Homalictus fijiensis* | 478.5 | 813 |
| *Homalictus groomi* | 904.0 | 108 |
| *Homalictus hadrander* | 828.8 | 66 |
| *Homalictus nadarivatu* | 963.4 | 113 |
| *Homalictus ostridorsum* | 1258.4 | 81 |
| *Homalictus taveuni* | 19.0 | 3 |

| | | |
|---|---|---|
| *Homalictus terminalis* | 1118.0 | 6 |
| *Homalictus tuiwawae* | 867.0 | 581 |
| *Homalictus new species* 1 | 638.0 | 1 |
| *Homalictus new species* 2 | 1068.0 | 2 |

**Table S3**. Mean pairwise phylogenetic distance (mpd.obs) for each mountain/coastal site, randomised mpd for each site (mpd.rand.mean), standardised effect of mpd vs randomised mpd (mpd.obs.z), and quantile (mpd.obs.p) comparison between observed and randomised site mpd. Phylogenetic diversity estimates for sites with only 1 taxa were not possible.

| Site | ntaxa | mpd.obs | mpd.rand.mean | mpd.rand.sd | mpd.obs.rank | mpd.obs.z | mpd.obs.p |
|---|---|---|---|---|---|---|---|
| 1 | 8 | 2538.16 | 2390.29 | 332.57 | 6252.50 | 0.44 | 0.63 |
| 2 | 4 | 1752.84 | 2391.44 | 549.49 | 1575.00 | -1.16 | 0.16 |
| 3 | 6 | 2784.51 | 2389.74 | 409.34 | 8004.00 | 0.96 | 0.80 |
| 4 | 5 | 2980.63 | 2386.66 | 465.10 | 9076.00 | 1.28 | 0.91 |
| 5 | 3 | 3121.18 | 2383.84 | 676.21 | 7808.50 | 1.09 | 0.78 |
| 6 | 9 | 2875.36 | 2393.63 | 304.79 | 9757.50 | 1.58 | 0.98 |
| 7 | 1 | - | - | - | - | - | - |
| 8 | 4 | 3105.03 | 2388.92 | 544.58 | 9358.50 | 1.31 | 0.94 |
| 9 | 3 | 2966.09 | 2382.96 | 676.80 | 6465.50 | 0.86 | 0.65 |
| 10 | 5 | 2946.71 | 2389.25 | 463.74 | 8929.00 | 1.20 | 0.89 |
| 11 | 13 | 2702.39 | 2390.92 | 221.13 | 9334.00 | 1.41 | 0.93 |
| 12 | 5 | 2946.71 | 2389.68 | 466.59 | 8908.50 | 1.19 | 0.89 |
| 13 | 1 | - | - | - | - | - | - |
| 14 | 1 | - | - | - | - | - | - |
| 15 | 1 | - | - | - | - | - | - |
| 16 | 3 | 3315.05 | 2386.98 | 675.63 | 9832.50 | 1.37 | 0.98 |
| 17 | 2 | 2617.14 | 2389.84 | 949.94 | 6899.00 | 0.24 | 0.69 |
| 18 | 1 | - | - | - | - | - | - |
| 19 | 1 | - | - | - | - | - | - |
| 20 | 1 | - | - | - | - | - | - |
| 21 | 1 | - | - | - | - | - | - |

| | | | | | | | |
|---|---|---|---|---|---|---|---|
| 22 | 1 | - | - | - | - | - | - |
| 23 | 2 | 3664.00 | 2364.08 | 942.12 | 7945.50 | 1.38 | 0.79 |
| 24 | 1 | - | - | - | - | - | - |
| 25 | 3 | 3008.10 | 2385.63 | 676.85 | 6812.50 | 0.92 | 0.68 |
| 26 | 6 | 2720.43 | 2388.52 | 411.60 | 7668.00 | 0.81 | 0.77 |
| 27 | 1 | - | - | - | - | - | - |
| 28 | 1 | - | - | - | - | - | - |
| 29 | 1 | - | - | - | - | - | - |

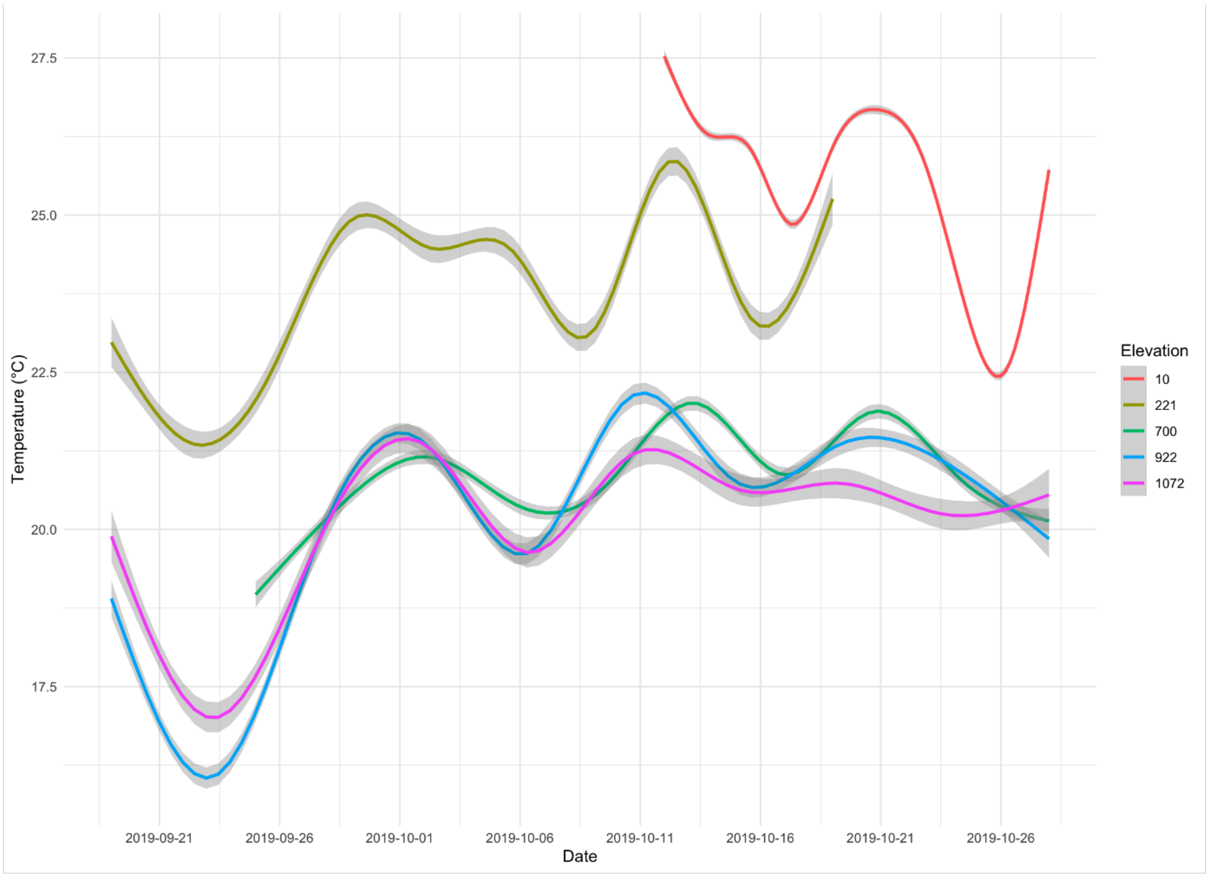


**Figure S1.** Environmental temperatures (from HOBO temperature loggers) at five sites across the elevational gradient from September/October 2019.

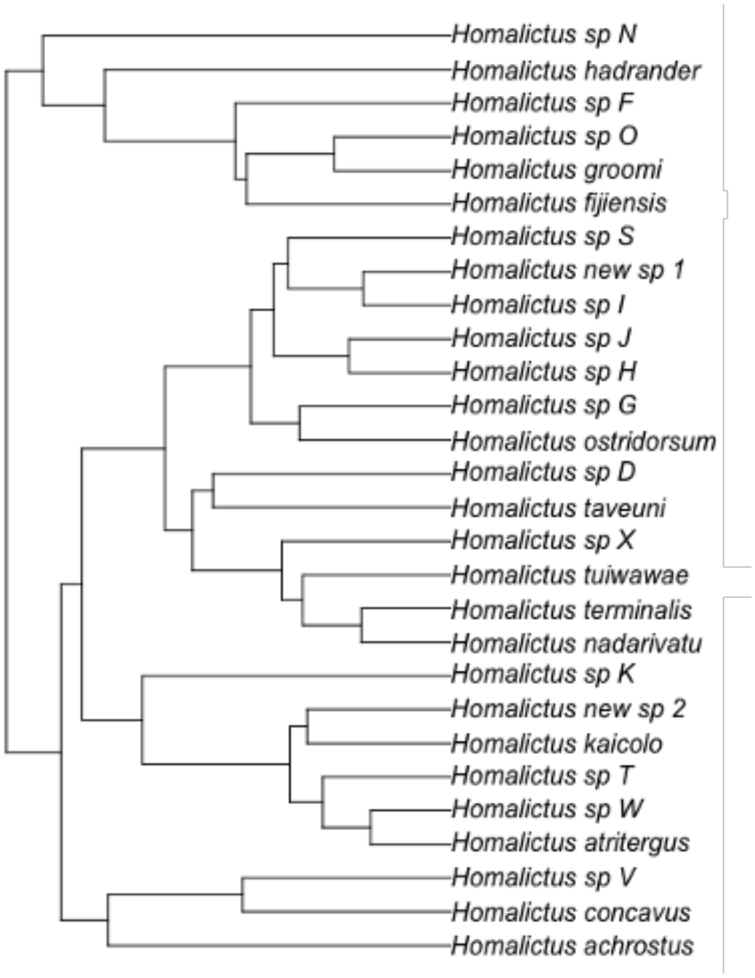


**Figure S2.** Phylogeny of all *Lasioglossum (Homalictus)* species (N = 28) sampled in Fiji between 2011 and 2019.

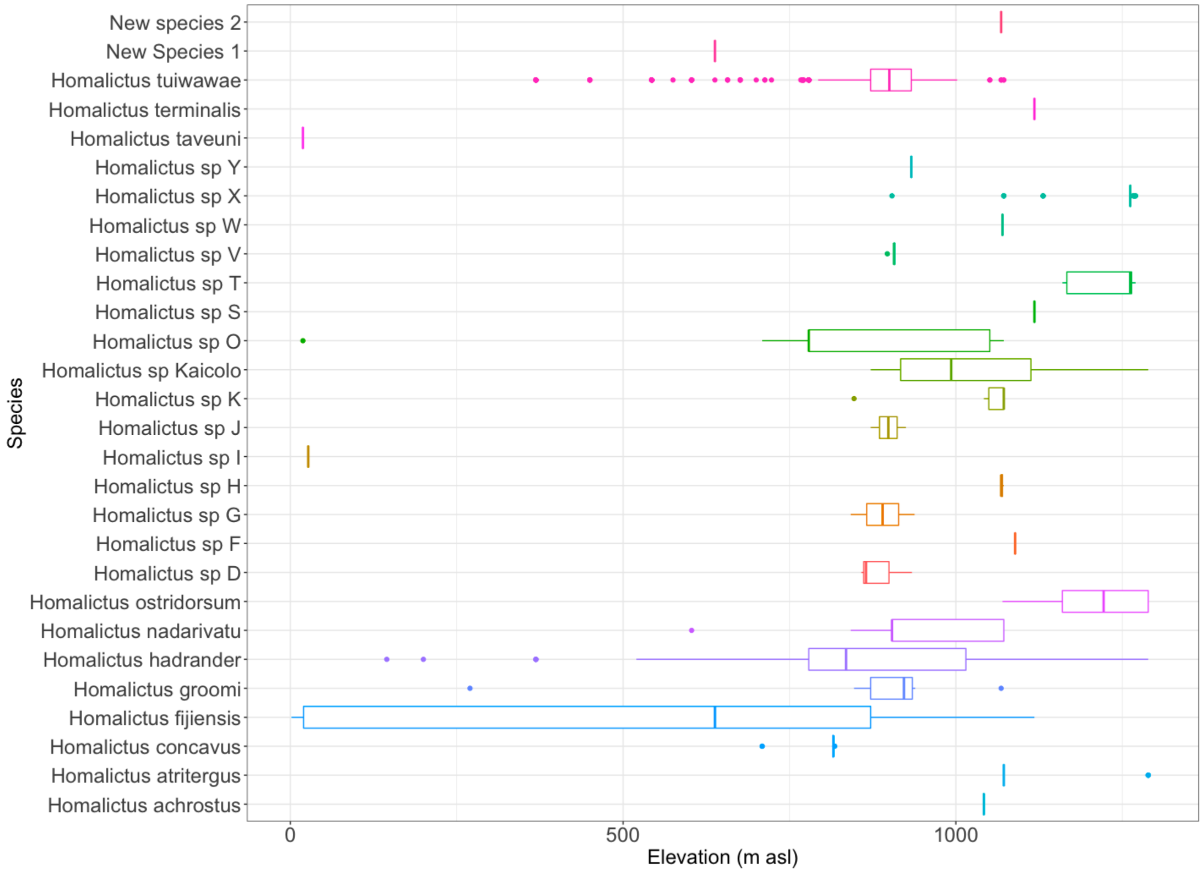


**Figure S3**. Elevational ranges of each *Lasioglossum (Homalictus)* species across the Fijian archipelago.
